# Cortical Complexity Alterations in Motor Subtypes of Parkinson’s Disease: A Surface-Based Morphometry Analysis of Fractal Dimension

**DOI:** 10.1101/2024.08.03.606491

**Authors:** Yousef Dehghan, Yashar Sarbaz

**Affiliations:** Biological Systems Modeling Laboratory, Department of Biomedical Engineering, Faculty of Electrical and Computer Engineering, University of Tabriz, Tabriz, Iran

**Keywords:** Parkinson’s disease, Tremor dominant, Postural instability gait difficulty, Structural MRI, Cortical folding

## Abstract

**Background:** Based on motor symptoms, Parkinson’s disease (PD) can be classified into tremor dominant (TD) and postural instability gait difficulty (PIGD) subtypes. This study aimed to investigate differences in cortical complexity and gray matter volume (GMV) between TD and PIGD.

**Methods:** We enrolled 36 TD patients, 27 PIGD patients, and 66 healthy controls (HC) from the PPMI (Parkinson’s Progression Markers Initiative) database. Voxel-based morphometry (VBM) and surface-based morphometry (SBM) were utilized to assess differences in GMV, cortical thickness, and cortical complexity. Additionally, correlations between clinical data and structural changes were examined.

**Results:** In comparison with HCs, PIGD patients exhibited a significant fractal dimension (FD) decrease in many cortical regions, such as the bilateral insula, right superior temporal, and left rostral middle frontal. Moreover, PIGD patients showed significant FD reduction in various regions, including the left supramarginal gyrus, left lateral orbitofrontal gyrus, right superior temporal gyrus, left lateral occipital gyrus, and bilateral insula, compared to the TD group. A significant negative correlation between age and FD was observed in the left insula for the PIGD patients and in the bilateral insula for the TD patients. However, no significant differences were found in GMV, cortical thickness, or other complexity indices.

**Conclusion:** Altered FD, particularly in bilateral insula, indicates that postural instability and gait disturbances in PD may result from a failure to integrate information from various structures, whereas parkinsonian tremor is not associated with this integration.

## 1. Introduction

Parkinson’s disease (PD) is one of the most common neurodegenerative disorders [1], which is characterized by the loss of dopaminergic neurons in the substantia nigra pars compacta and the aberrant accumulation of a-synuclein in brain tissue [2]. The main clinical manifestations of PD include rest tremor, bradykinesia, rigidity, and postural instability [3]. Prior studies have demonstrated that PD can be categorized into various subtypes based on differences in clinical symptoms, disease development, prognosis, and responsiveness to treatment [4–6]. The three most common PD subtypes are postural instability gait difficulty (PIGD), tremor dominant (TD), and indeterminate or mixed. The patients in the intermediate group manifest both TD and PIGD subtype symptoms to an almost equal extent. The TD patients have a better outlook, whereas the PIGD patients have increased cognitive impairment and do not respond well to dopamine replacement therapy [7–9].

Magnetic resonance imaging (MRI) has been extensively used for diagnosing PD and other neurodegenerative diseases [10]. Voxel-based morphometry (VBM) is a method for comparing 3-D brain scans voxel-by-voxel across different subject groups in order to calculate differences within regional gray matter concentrations [11, 12]. Surface-based morphometry (SBM) is an additional methodology utilized to estimate cortical morphology based on structural MRI [13, 14]. While SBM assesses a variety of features, such as cortical thickness, fractal dimension, gyrification index, and sulcal depth (the latter three represent cortical complexity) [15], VBM estimates differences in volumetric features like gray matter volume (GMV). The gyrification index (GI) is expressed as the ratio of the inner surface area to the outer surface area of an outer convex hull [16]. Sulcal depth represents the quantification of the distance toward an idealized smooth brain surface [17], explicitly measuring the depth of the sulci within the brain. According to Yotter *et al*. [16], the localized measurement of fractal dimension (FD) can be calculated using spherical harmonic reconstructions. Also, FD can be noticed as a method of approximating gyrification by considering parameters such as sulcal depth, convolution of gyral pattern, and cortical folding frequency [18]. Utilizing both VBM and SBM in the analysis can be advantageous in identifying subtle morphological alterations, particularly in the cortex, that are caused by psychiatric or neurological disorders [19].

Many studies have investigated morphological differences between PD motor subtypes. Most previous studies have investigated possible volumetric or thickness alterations. One instance is a specific study that examined GMV differences between predominate PIGD and predominate TD patients without comparing any of these groups to a control group [20]. The study found decreased GMV across all primary brain lobes and subcortical regions in PIGD compared to TD. Cerasa *et al*. [21] indicated increased GMV in the bilateral inferior frontal gyrus in dyskinetic PD compared to non-dyskinetic PD. Zheng *et al*. [22] reported increased GMV in the medial frontal gyrus in the TD group compared to PIGD. While other studies did not find any differences in the volume or structure of the cortex between TD and PIGD [23–26], Herb *et al*. [27] demonstrated that the PIGD patients have decreased cortical thickness in the dorsolateral frontal lobes, anterior temporal lobes, cuneus, and precuneus compared to the TD patients. Li *et al*. [28] investigated cortical and volumetric subcortical changes between TD, akinetic-rigid, and a control group. In comparison to the HC group, the study found increased sulcal depth in the right supramarginal gyrus in the TD subtype and a volumetric decrease in the hippocampus of both subtypes. However, there were no significant differences between the two subtypes in terms of sulcal depth or subcortical volume changes.

FD has been extensively used to investigate alterations in the morphological cerebral pattern in various conditions and neurological disorders, such as amyotrophic lateral sclerosis [29], minimal hepatic encephalopathy [30], anorexia nervosa [31], multiple sclerosis [32], spinocerebellar ataxia [33], Alzheimer’s disease [34], and dementia spectrum disorders [35]. There are a few studies that have investigated FD changes in PD patients. In a cross-sectional study, Li *et al*. [36] investigated changes in cortical FD and gyrification between non-demented PD and HCs. The study found widespread FD reductions in PD patients compared to HCs but no significant differences in GI. Zhang *et al*. [37] studied differences in cortical complexity between early-stage right-onset and left-onset PD and HCs. They found that right-onset PD patients had decreased FD in the left superior temporal cortex compared to HCs. Yuan *et al*. [38] investigated volumetric and cortical differences between PD patients with and without depression and an HC group. The study indicated decreased GI in depressed PD patients compared to non-depressed PD and HCs. However, the difference in FD or sulcal depth was not significant. Also, widespread cortical atrophy was observed in depressed PD patients.

Previous studies, however, have not yet adequately assessed differences in cortical surface complexity (FD, GI, and sulcal depth) in PD patients with different motor symptoms. Furthermore, studies focusing on GMV, or cortical thickness, reported inconsistent results. In the present study, we aimed to investigate the changes in both cortical characteristics and subcortical structures among PIGD, TD, and a sex- and age-matched control group. We enhanced our SBM analysis by incorporating the VBM analysis to identify potential cortical differences (FD, GI, sulcal depth, and thickness) as well as volumetric alterations.

## 2. Materials and methods

In this study, we utilized a combination of VBM and SBM in our analysis, with a particular emphasis on the cortical regional FD. We described the statistical tests for the demographic data of the subjects in the participants section and the voxel-wise and vertex-wise statistical tests in the data Processing section. Also, the summarized workflow of the processing pipeline is illustrated in Figure 1.

**Figure 1.**
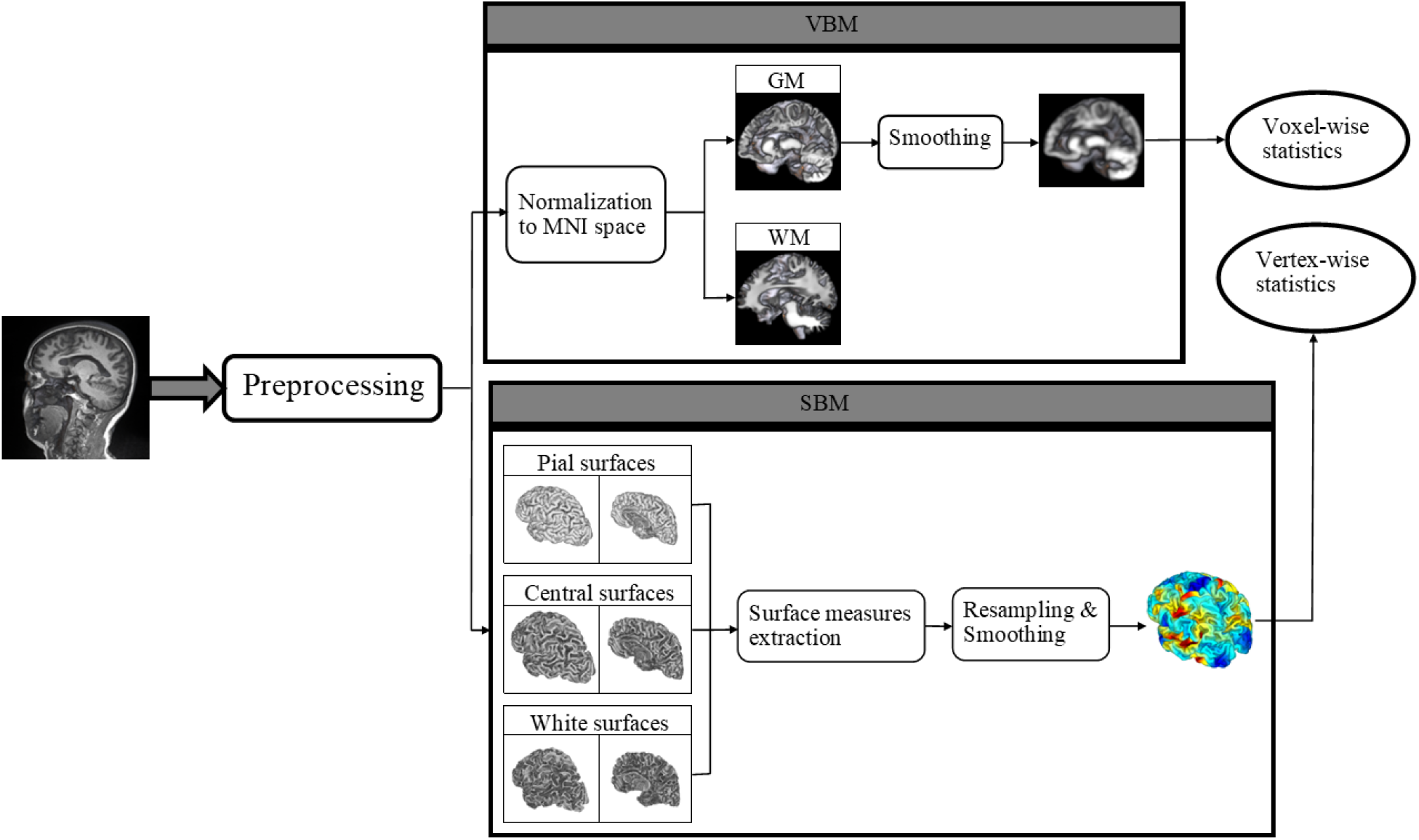
The workflow of the processing pipeline. GM, gray matter; WM, white matter

### 2.1. Participants

This study included 168 PD patients and 66 healthy controls (HC). All MRI and clinical data were obtained on March 15, 2024, from the Parkinson’s Progression Markers Initiative (PPMI) database (https://www.ppmi-info.org/access-data-specimens/download-data), a multicenter longitudinal observational study of PD [39]. Since this study is based on an open-source database, written informed consent was waived. We included only the data acquired during the baseline visit for all subjects. Moreover, enrolled PD patients are required to be untreated with levodopa or other PD treatments.

Based on MDS-UPDRS (MDS-UPDRS TD/PIGD ratio), PD patients were classified into three groups: PIGD (ratio less than 0.9), TD (ratio greater than 1.15), and intermediate (ratio ≤ 1.15 and ≥ 0.9) [40]. To reduce confounding factors and improve detection of subtle structural differences between the TD and PIGD subtypes, we excluded 105 patients from the study. The exclusion criteria were as follows: 1) patients categorized into the intermediate group; and 2) TD patients with a TD/PIGD ratio less than 3. We made this decision to reduce heterogeneity within the study groups and increase the likelihood of detecting alterations specific to each subtype of the disease in between-group comparisons.

The statistical analyses of demographic and clinical characteristics were conducted using a chi-square test for gender, an analysis of variance (ANOVA) for age, and the Mann-Whitney U test for MDS-IPDRS III, the H&Y stage, tremor score, and PIGD score.

### 2.2. MRI data acquisition

Structural MRIs were collected using a 3-T Siemens Trio Trim system. The acquisition parameters of the 3D T1-weighted images were as follows: TR/TE/TI = 2300/3/900 ms, flip angle = 9°, slice thickness = 1.00 mm, voxel size = 1.0 × 1.0 × 1.0 mm^3^, and sagittal orientation with a 256 × 256 × 176 matrix size.

### 2.3. Data processing

All imaging data were processed and statistically analyzed using CAT12 (https://neuro-jena.github.io/cat/) implemented in SPM12 (https://www.fil.ion.ucl.ac.uk/spm/software/spm12/).

#### 2.3.1. Preprocessing

We preprocessed the 3D T1-weighted MRI data using the following key steps: First, we applied a spatially adaptive non-local means filter [41] to reduce the noise. Subsequently, all images were bias-corrected and affine-registered to a prior tissue probability map, CAT12’s default probability map. The “Unified Segmentation” approach [42] was utilized for the initial estimation of segmented tissue classes but not for the final segmentation. After stripping the skull, the brain was parcellated into subcortical regions and two discrete hemispheres, followed by the local intensity correction performed for all tissue classes. Finally, an adaptive maximum a posteriori method [43], which is independent of prior tissue probabilities, was used for the segmentation of the brain into three tissue classes: gray matter (GM), white matter (WM), and cerebrospinal fluid (CSF). The final segmentation was refined with a partial volume estimation [44] to reduce the partial volume effects and increase the segmentation accuracy for the voxels with mixed tissue types.

#### 2.3.2. Voxel-based analysis

The segmented GM was spatially normalized to Montreal Neurological Institute 152 (MNI152) space using geodesic shooting registration [45]. Then, all normalized GM volumes were smoothed using a Gaussian kernel with an 8-mm full width at half maximum (FWHM).

The CAT12 statistical module was used to perform an ANOVA and t-tests to identify potential group-level differences in GMV. Age, sex, and total intracranial volume were included as covariates in the statistical tests mentioned above. The resulting parametric statistical maps were corrected for multiple comparisons, with statistical significance defined as *p* < 0.05.

#### 2.3.3. Surface-based analysis

First, the initial cortical thickness and the initial central surface of both hemispheres were estimated [15]. After the correction of topological defects [46], the following final surface meshes were obtained for each hemisphere separately: pial, white, and central. Consequently, the central surface meshes were reparametrized and spatially registered [16] to the Freesurfer “FsAverage” template. Cortical thickness was computed using the projection-based thickness (PBT) method [15]. The PBT method allowed for estimation of the thickness and reconstruction of the central surface, both at the same step. Moreover, GI, FD, and sulcal depth were extracted from the central surfaces of the left and right hemispheres.

For FD estimation, the spherical harmonic coefficients of central surfaces must be extracted to analyze the harmonic content of re-parameterized spherical meshes using 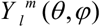 :

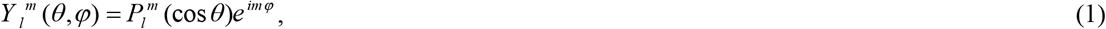

Where *θ* and *φ* are the spherical coordinates (*θ* is the co-latitude and *φ* is the azimuthal coordinate). Also, *m* and *l* are integers, and 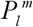 is the function defined by:

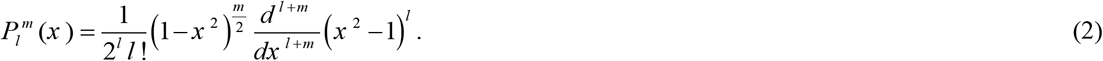

The function *f* (*θ, φ*) that is square-integrable on the sphere can be determined by

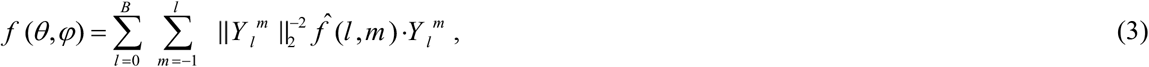

Where *B* is the maximum *l*-value. Each 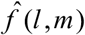, as a coefficient, is defined by 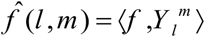, and the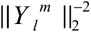 is calculated by:

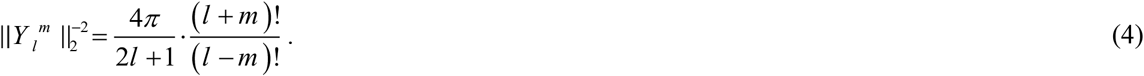

After obtaining the coefficients, ten surfaces were reconstructed from each central surface, and local FD can be calculated by finding the slope of a regression plot at vertex level: log (area) versus log (the maximum *l*-value of the reconstruction) [16]. The local GI was obtained on the basis of absolute mean curvature values [47]. The mean curvature at vertex level is determined as

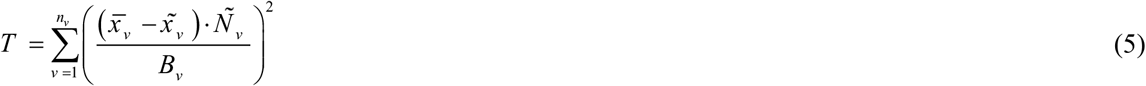

where 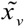is the centroid of the neighboring vertices of vertex *v*, and *B*_*v*_ denotes the average distance from each neighboring vertex to this centroid. Also, the Euclidean distance between the central surface and its convex hull was utilized to calculate the sulcal depth. Finally, the surface measures from the left and right hemispheres were merged, resampled, and smoothed using a Gaussian kernel with a 15-mm FWHM for thickness and a 20-mm FWHM for FD, GI, and sulcal depth.

For the statistical analyses, we did an ANOVA and t-tests with the CAT12 statistical module to see if there were any group-level differences for each of the mentioned surface-based morphometric measures. Age and sex were defined as covariates in the mentioned statistical test. The parametric maps resulting from the mentioned statistical tests were enhanced by employing threshold-free cluster enhancement (TFCE) [48], which was implemented in CAT12. Statistical significance was defined as *p* < 0.05, and results were corrected for multiple comparisons using the false discovery rate [49]. After defining the regions with significant differences in each surface measure according to the Desikan-Killiany atlas [50], the regional surface measures were extracted from the clusters with significant differences in between-group comparisons. Finally, Spearman’s correlation coefficient was used to perform a correlational analysis between the raw surface data extracted from significant clusters and the clinical data of participants, defining statistical significance as *p* < 0.05 (Holm-Bonferroni corrected).

## 3. Results

### 3.1. Descriptive analysis

A total of 36 TD patients, 27 PIGD patients, and 66 matched controls were ultimately included in the analysis. The demographic and clinical data of the participants are shown in Table 1. No significant differences were seen in age, gender, Hoehn & Yahr stage, or UPDRS III score. Nevertheless, the tremor scores in the TD group were significantly higher than in the PIGD group. Furthermore, the PIGD scores were significantly higher in the PIGD group compared to the TD group.

**Table 1.**
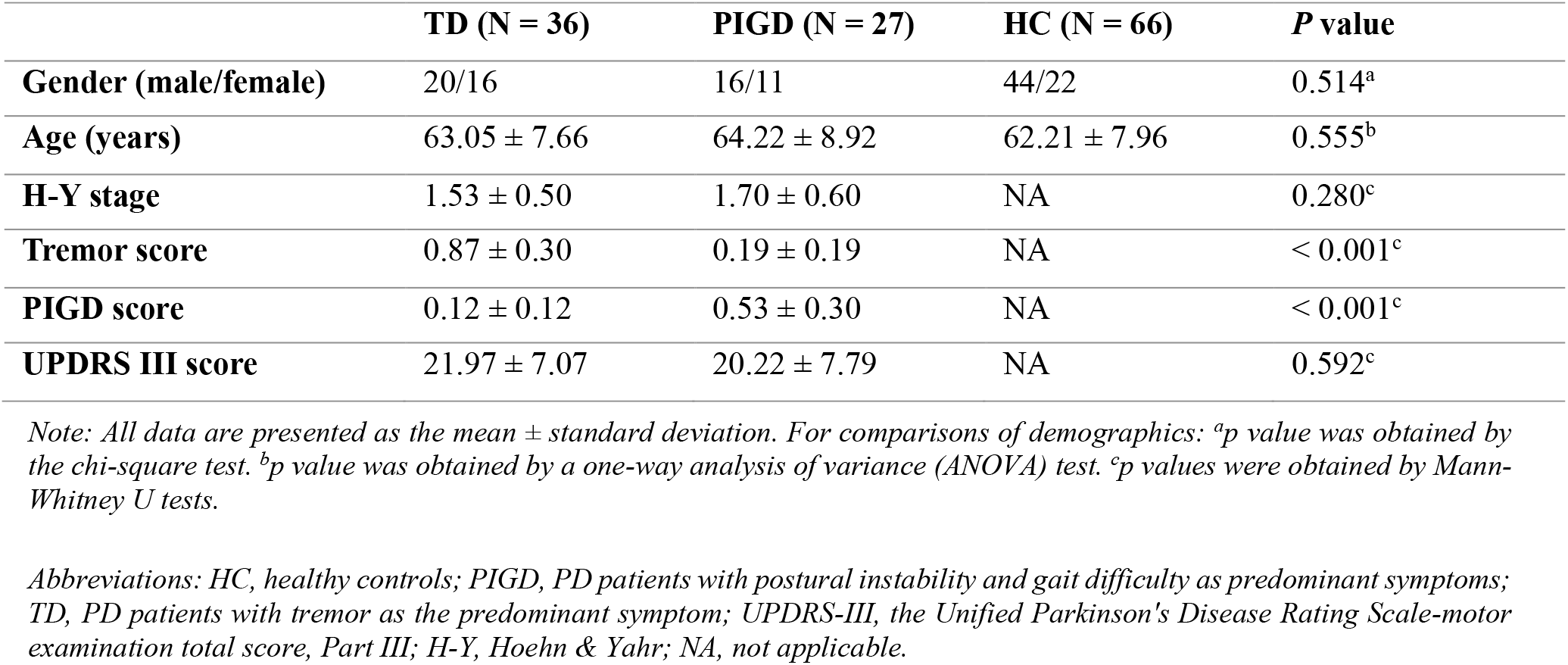
The demographics and clinical data of all participants.

|  | TD (N = 36) | PIGD (N = 27) | HC (N = 66) | P value |
| --- | --- | --- | --- | --- |
| **Gender (male/female)** | 20/16 | 16/11 | 44/22 | 0.514<sup>a</sup> |
| **Age (years)** | 63.05 ± 7.66 | 64.22 ± 8.92 | 62.21 ± 7.96 | 0.555<sup>b</sup> |
| **H-Y stage** | 1.53 ± 0.50 | 1.70 ± 0.60 | NA | 0.280<sup>c</sup> |
| **Tremor score** | 0.87 ± 0.30 | 0.19 ± 0.19 | NA | < 0.001<sup>c</sup> |
| **PIGD score** | 0.12 ± 0.12 | 0.53 ± 0.30 | NA | < 0.001<sup>c</sup> |
| **UPDRS III score** | 21.97 ± 7.07 | 20.22 ± 7.79 | NA | 0.592<sup>c</sup> |

Note: All data are presented as the mean ± standard deviation. For comparisons of demographics: <sup>a</sup>*p* value was obtained by the chi-square test. <sup>b</sup>*p* value was obtained by a one-way analysis of variance (ANOVA) test. <sup>c</sup>*p* values were obtained by Mann-Whitney *U* tests.
Abbreviations: HC, healthy controls; PIGD, PD patients with postural instability and gait difficulty as predominant symptoms; TD, PD patients with tremor as the predominant symptom; UPDRS-III, the Unified Parkinson's Disease Rating Scale-motor examination total score, Part III; H-Y, Hoehn & Yahr; NA, not applicable.

### 3.2. Gray matter volume and cortical thickness analysis

VBM results are provided in Table 2. According to this table, no significant differences in GMV were found between the PIGD and HC groups, nor between the TD and HC groups. Furthermore, a comparison between the PIGD and TD groups revealed no significant differences in GMV. Additionally, the between-group comparisons did not reveal any significant differences in cortical thickness. Also, ANOVA revealed no significant differences in GMV or cortical thickness among the TD, PIGD, and HC groups.

**Table 2.** VBM results.

|  | Cluster size | *p*-value<br>(uncorrected) | *p*-value<br>(FWE<br>corrected) | Peak MNI coordinates |  |  |
| --- | --- | --- | --- | --- | --- | --- |
|  |  |  |  | X | Y | Z |
| **Regions of GMV reduction**<br>**between PIGD and HC** | 11 | 0.693 | 0.999 | 8 | -82 | 2 |
|  | 9 | 0.725 | 0.999 | 12 | -69 | -2 |
| **Regions of GMV reduction**<br>**between TD and HC** | 94 | 0.209 | 0.870 | -36 | -14 | -34 |
|  | 66 | 0.290 | 0.941 | -20 | 15 | 70 |
|  | 30 | 0.481 | 0.991 | -4 | 22 | 68 |
|  | 30 | 0.481 | 0.991 | -26 | -10 | 66 |
|  | 18 | 0.595 | 0.997 | 48 | -22 | -34 |
|  | 14 | 0.644 | 0.998 | -50 | -6 | -20 |
|  | 11 | 0.688 | 0.999 | 20 | -69 | 64 |

Abbreviations: HC, healthy controls; PIGD, PD patients with postural instability and gait difficulty as predominant symptoms; TD, PD patients with tremor as the predominant symptom; GMV, gray matter volume.

### 3.3. Cortical complexity analysis

PIGD patients have demonstrated widespread decreases in regional FD when compared to both the HC group and the TD patients. Comparing PIGD patients and HCs, the FD analysis revealed significant clusters in both the left and right hemispheres. The cortical regions with a significant difference in FD between PIGD patients and HCs were illustrated in Figure 2a, and detailed information about clusters is shown in Table 3. According to table 3, the largest cluster in the right hemisphere is composed of 1451 vertices (*p* = 0.01476, FDR corrected after TFCE). This cluster is located in the insula, superior temporal gyrus, supramarginal gyrus, postcentral gyrus, precentral gyrus, middle temporal gyrus, and transverse temporal gyrus. Additionally, the largest cluster in the left hemisphere is composed of 3047 vertices (*p* = 0.01476, FDR corrected after TFCE), which is located in the insula, rostral middle frontal gyrus, lateral orbitofrontal gyrus, supramarginal gyrus, superior frontal gyrus, precentral gyrus, caudal middle frontal gyrus, pars opercularis, postcentral gyrus, pars triangularis, and transverse temporal gyrus.

**Table 3.** Regions of decreased fractal dimension in Parkinson’s disease with postural instability and gait difficulty (PIGD) compared to healthy controls (HC) after threshold-free cluster enhancement (p < 0.05, FDR corrected).

| Hemisphere | Cluster size | p-value | Overlap of atlas region |
| --- | --- | --- | --- |
| LH | 3047 | 0.01476 | 23% insula<br>17% rostral middle frontal<br>13% lateral orbitofrontal<br>11% supramarginal<br>8% superior frontal<br>8% precentral<br>7% pars opercularis<br>7% caudal middle frontal<br>2% postcentral<br>2% pars triangularis<br>1% transverse temporal |
| RH | 1451 | 0.01476 | 48% insula<br>32% superior temporal<br>9% supramarginal<br>6% postcentral<br>2% precentral<br>1% middle temporal<br>1% transverse temporal |
| LH | 68 | 0.03070 | 100% paracentral |
| LH | 58 | 0.04002 | 100% rostral middle frontal |
| LH | 6 | 0.04851 | 50% lingual<br>50% precuneus |
| RH | 5 | 0.04851 | 60% insula<br>20% lateral orbitofrontal |
| LH | 1 | 0.04851 | 100% rostral middle frontal |

Atlas labeling was performed according to the Desikan-Killiany atlas.
Abbreviations: LH, left hemisphere; RH, right hemisphere.

**Figure 2.**
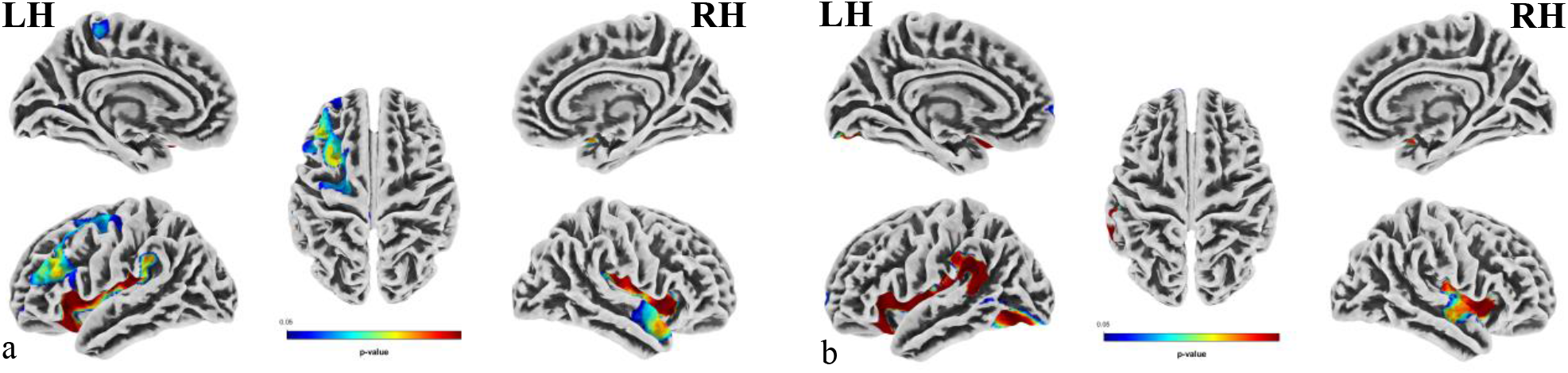
a. Fractal dimension (FD) reduction regions in Parkinson’s disease with postural instability gait difficulty (PIGD) compared to healthy controls (HC) (P < 0.05, FDR corrected after TFCE). b. Fractal dimension (FD) reduction regions in Parkinson’s disease with postural instability gait difficulty (PIGD) compared to tremor dominant (TD) subtype of the Parkinson’s disease (P < 0.05, FDR corrected after TFCE).

Furthermore, in comparison between PIGD and TD patients, the FD analysis revealed significant clusters. Figure 2b illustrates the cortical regions with a significant difference in FD between PIGD and HC patients, and Table 4 provides detailed information about clusters. According to table 4, in the right hemisphere, the significant cluster consisted of 1106 vertices (p = 0.02812, FDR corrected after TFCE) and was located in the insula, superior temporal gyrus, and transverse temporal gyrus. The largest cluster in the left hemisphere is composed of 2182 vertices (p = 0.02812, FDR corrected after TFCE), which is located in the insula, supramarginal gyrus, lateral orbitofrontal gyrus, superior temporal gyrus, banks of superior temporal sulcus, transverse temporal gyrus, postcentral gyrus, and medial orbitofrontal gyrus. However, PIGD patients did not have increased FD in any cortical region. Furthermore, we found no significant differences in FD between TD patients and HCs. ANOVA also revealed no statistically significant differences among the TD, PIGD, and HC groups. Notably, we observed no significant differences in GI or sulcal depth among the PIGD, TD, and HC groups.

**Table 4.** Regions of decreased fractal dimension in Parkinson’s disease with postural instability and gait difficulty (PIGD) compared to tremor-dominant (TD) subtype of the Parkinson’s disease after threshold-free cluster enhancement (p < 0.05, FDR corrected).

| Hemisphere | Cluster size | *p*-value | Overlap of atlas region |
| --- | --- | --- | --- |
| LH | 2182 | 0.02812 | 32% insula<br>26% supramarginal<br>20% lateral orbitofrontal<br>9% superior temporal<br>6% banks sts<br>3% transverse temporal<br>2% postcentral<br>1% medial orbitofrontal |
| RH | 1106 | 0.02812 | 48% superior temporal<br>45% insula<br>6% transverse temporal |
| LH | 435 | 0.02812 | 57% lateral occipital<br>31% fusiform<br>11% inferior temporal |
| LH | 31 | 0.04208 | 97% superior frontal<br>3% rostral middle frontal |
| LH | 22 | 0.04342 | 64% middle temporal<br>23% inferior temporal<br>14% lateral occipital |
| LH | 10 | 0.04463 | 100% middle temporal |
| LH | 6 | 0.04725 | 100% lateral occipital |

*Atlas labeling was performed according to the Desikan-Killiany atlas.*
*Abbreviations: LH, left hemisphere; RH, right hemisphere.*

### 3.4. Correlation analysis

In TD patients, we found a significant negative correlation between age and regional FD of the right insula (*r* = -0.588, *p* = 0.003, Holm-Bonferroni corrected) and the left insula (*r* = -0.631, *p* = 0.0007, Holm-Bonferroni corrected). Furthermore, in the PIGD group, we observed a significant negative correlation between age and the regional FD of the left insula (*r* = -0.59, *p* = 0.0284, Holm-Bonferroni corrected). Figure 3 illustrates the scatter plots of these associations. However, no significant correlation between regional FD of other cortical regions and clinical characteristics was found during the analysis.

**Figure 3.**
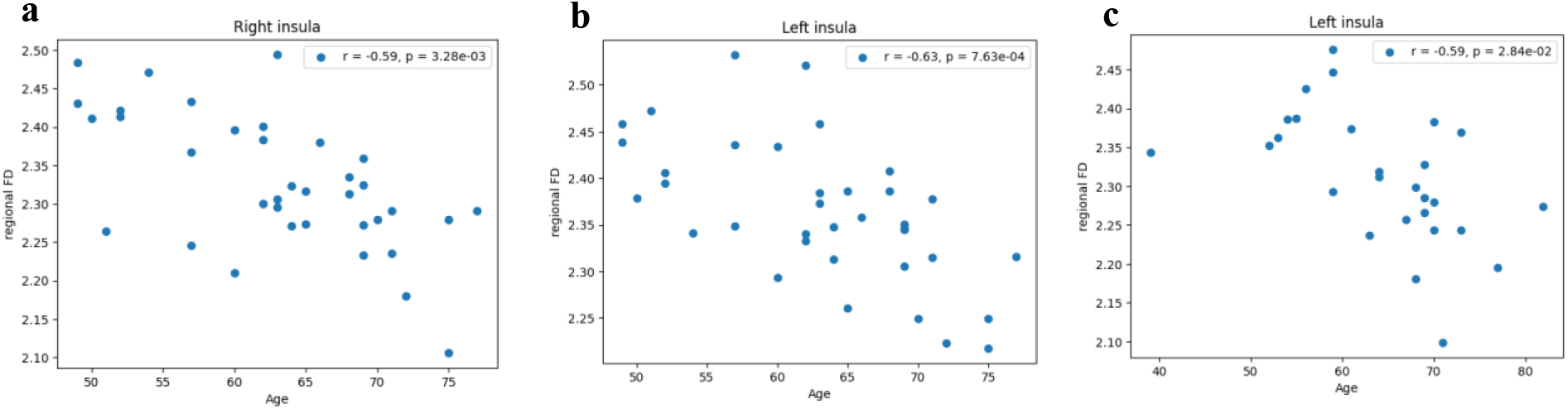
a. scatterplot representing the negative correlation between regional FD of the right insula and age in the TD group. b. scatterplot representing the negative correlation between regional FD of the left insula and age in the TD group. c. scatterplot representing the negative correlation between regional FD of the left insula and age in the PIGD group.

## 4. Discussion

In this study, we investigated the differences in cortical surface complexity between two distinct subtypes of PD patients with different motor symptoms in the early stages of the disease and healthy participants. Additionally, we assessed the differences in cortical atrophy patterns and subcortical volumetric alterations in each subtype. Compared to HC participants, PIGD patients exhibited bilateral decreased FD in all main cortical brain lobes except the occipital lobe. Furthermore, comparing PIGD patients to the TD group, we observed FD reductions in all left-hemisphere cortical lobes but only in the temporal and insular lobes of the right hemisphere. However, we found no significant differences in FD between TD and HC. Our analysis revealed no significant differences in GMV or cortical atrophy between these patient groups and HC.

The nervous system has developed a fractal structure [51, 52]. Also, it is posited that the cerebral cortex has fractal characteristics [53]. However, the cerebral cortex is distinguished by a heightened complexity relative to other components of the human nervous system. Both FD and GI are quantifications of the cortical folding pattern. While high FD or GI in a specific cortex region indicates increased cortex folding, lower FD or GI indicates a reduction of cortical folding. Neurodegenerative diseases can induce alterations in this fractal structure, reducing the fractal dimension in various regions of the cortex [54]. This alteration in the cortical fractal structure may be an indirect consequence of damage sustained by the brain’s subcortical regions. Previous studies noted altered cortical complexity in PD patients [36–38]. Clinically, the symptoms manifested by PD patients could be heterogeneous [3]. It is essential to acknowledge that the symptomatology in PD patients is not uniform, and one group of symptoms can be presented as predominant during the disease’s incipient phase. Conversely, a full spectrum of symptoms is typically observed in the disease’s advanced stages [55].

In the current study, the most robust finding was a decreased FD of bilateral insula in the PIGD group, compared to both the HC and TD groups. As an anatomical integration hub, the insular cortex contributes to processing the sensory information received from various cortical and subcortical regions, emotion regulation, and cognitive functions [56]. Previous studies suggested increased cognitive impairments in PIGD rather than TD [7, 9]. The role of the insular cortex in emotional processing and cognitive function suggests a possible relationship between FD reduction in the insular cortex and higher cognitive impairment in PIGD patients. The insula, with its connections to the striatum and its role in integrating sensory and motor information, plays a crucial role in coordinating movements and maintaining balance [57]. Therefore, we hypothesize that the decreased FD in the insular region may be a reflection of damage to connections between the insula and basal ganglia or other subcortical structures, leading to a failure in this region’s integrative function. However, FD reductions in multiple cortical regions may serve as an indicator for slightly altered fiber tracts in deep white matter, especially in the initial stages of the disease.

In the PIGD group, we found a statistically significant negative correlation between age and the regional FD of the left insula. Also, the regional FD of the bilateral insula was negatively correlated with age in the TD group. Given that this correlation was observed in both subtypes, disease progression with advancing age is the most plausible explanation for this finding. This association could indicate that age-related structural changes of the insula in PD patients are more common compared to other cortical regions, which could be related to the integrative functions of this structure.

Our analysis revealed no significant differences in the cortical FD of the TD subtype compared to the HC group. The presence of a tremor itself is not a definitive diagnostic criterion for PD [9, 58]. Additionally, tremors can be attributed to various etiologies, including dystonic, essential, or psychogenic origins, and are not solely indicative of neurodegeneration. Tremor manifestations may result from dysregulation within the motor control system and are not exclusively due to acute cerebral trauma [59]. Consequently, individuals with TD Parkinsonism have relatively preserved balance control capabilities, in contrast to those with PIGD, who show more significant structural brain damage due to impaired balance regulation. The absence of significant differences in cortical FD between TD patients and HCs highlights the regulatory, rather than structural, nature of their condition. On the other hand, PIGD patients who have trouble controlling their balance have structural damage to the brain that is severe enough to reduce the FD across multiple cortical regions.

Our findings on reduced FD, but not other complexity measures, align with the results of Li *et al*. [36]. However, in contrast to the current study, they reported FD reductions in different cortical regions. The following arguments can potentially address this discrepancy in the results: (1) We limited our study to patients who were in the initial stages of the disease and had not received any treatment yet; (2) TD patients with low tremor scores were excluded from this study. PD patients manifest a vast spectrum of clinical symptoms, and most of the patients can be categorized into the excluded group.

According to Yuan *et al*. [38], the GI of the bilateral insular cortex has significantly decreased in PD with depression. Additionally, Zhang *et al*. [60] reported that PD patients demonstrated altered GI in various cortical regions, such as the lingual and fusiform gyri, the orbitofrontal cortex, and the temporal gyri. In contrast, our study identified decreased FD in the bilateral insula and several key regions highlighted by Zhang *et al*. [60] rather than changes in GI. By situating our findings within the context of previous research, we speculate that FD may serve as a more sensitive measure than GI, potentially predicting cortical changes in the later stages of the disease, especially in the PIGD subtype.

The basal ganglia’s dysfunction has a crucial role in the onset of tremor symptoms in PD [61]. PIGD patients exhibit various FD alterations in cortical regions such as the insula, implying that abnormalities in the integrative system are associated with postural instability and gait disturbances. This suggests that these symptoms stem from a failure to integrate information from different cortical and subcortical regions effectively. The TD subtype, on the other hand, doesn’t show any changes in cortical FD, especially in the insular cortex. This implies that a malfunction in a subcortical structure such as the basal ganglia could be the cause of parkinsonian tremor, rather than a failure in the integrative system.

However, there are several limitations in our study that need to be acknowledged. First, as cross-sectional research, only the data acquired during the baseline visit was included in our study. It is clear that PD patients exhibit different symptoms as the disease progresses and at different stages. A patient with a TD subtype in the initial stage may develop severe instabilities in later stages of the disease, leading to categorization into other subtypes such as PIGD or the intermediate group. So, additional longitudinal studies will be needed to assess the time-related changes in cortical characteristics. Another limitation of our study is the exclusion of patients with low tremor scores from the TD group due to a reduction in heterogeneity within this cohort. Furthermore, to mitigate the confounding effects of dopaminergic therapies, our investigation has exclusively examined patients without pharmacological PD treatment. However, additional studies are warranted to determine how much the consumed medications influence the degree of alterations in cortical features, such as the cortical FD, in each subtype.

## 5. Conclusion

The present study explored the volumetric and cortical differences between PD motor subtypes and a HC group. Our findings, which show altered FD in various cortical regions, including bilateral insula, in the PIGD group but not in the TD subtype, may define postural instability and gait disturbances as improper integration of information from different cortical and subcortical regions, while tremor is a malfunction of subcortical regions rather than an integration problem. Also, decreased FD in the PIGD subtype suggests that FD may serve as a novel biomarker in studying alterations in cortical morphology patterns or the diagnosis of the PIGD subtype. Moreover, widespread reductions in FD, but not in GI or sulcal depth, indicate that cortical FD is more sensitive than other measures of cortical complexity and possibly indicate information about structural changes that will happen in the later stages of the disease as it advances. Our results provide morphological support for the presence of differences in cortical complexity between the TD and PIGD subtypes. Our findings also suggest that the differences in GMV or cortical atrophy patterns reported in previous studies may be false positive results or depend on the later stages of the disease.

